# Reward responses to vicarious feeding depend on BMI

**DOI:** 10.1101/2024.06.20.599673

**Authors:** Lili Järvinen, Severi Santavirta, Vesa Putkinen, Henry K. Karlsson, Kerttu Seppälä, Lihua Sun, Matthew Hudson, Jussi Hirvonen, Pirjo Nuutila, Lauri Nummenmaa

**Author notes:** **Address Correspondence to:** Lili Järvinen, Turku PET Centre c/o Turku University Hospital FI-20520 Turku, Finland.

## Abstract

Eating is inherently social for humans. Yet, most neuroimaging studies of appetite and food-induced reward have focused on studying brain responses to food intake or viewing pictures of food alone. Here we used functional magnetic resonance imaging (fMRI) to measure haemodynamic responses to “vicarious” feeding. The subjects (n=97) viewed series of short videos representing naturalistic episodes of social eating intermixed with videos without feeding / appetite related content. Viewing the vicarious feeding (versus control) videos activated motor and premotor cortices, thalamus, and dorsolateral prefrontal cortices, consistent with somatomotor and affective engagement. Responses to the feeding videos were also downregulated as a function of the participants’ BMI. Altogether these results suggest that seeing others eating engages the corresponding motor and affective programs in the viewers’ brain, potentially increasing appetite and promoting mutual feeding.

## Introduction

Eating is inherently social for humans. Our species must feed their offspring since birth to ensure their survival, but the social nature of feeding extends all the way into adulthood. Every day, families, friends, and coworkers gather around breakfasts, dinners, and suppers, and it is almost impossible to think about human festivities without shared drinks and meals. Food sharing across others has likely evolved from sharing with offspring and partners to support coalitions and mate choice (Jaeggi & Van Schaik, 2011). The benefits of shared meals in the family after childhood are also present today, as eating together with the family is associated with better behavior and mental health as well as less substance abuse and suicidality in teenagers (Eisenberg et al., 2004, 2008; Meier, n.d.).

Social eating has also its downsides. For example, people eat larger portions when they are eating together than alone (Higgs & Thomas, 2016; Ruddock et al., 2021), possibly because of longer meals due to the social contact (Hetherington et al., 2006). Eating together, especially unhealthy food, is also more rewarding which may increase the intake of unhealthy food (Huang et al., 2022). Overall, social component of eating has been supposed to be a contributory factor of development and maintaining of obesity (Higgs & Thomas, 2016). Additionally, there is significant non-genetic social component to development of obesity, underlining the social transmission of unhealthy eating habits in social networks (Christakis & Fowler, 2007).

During the last decade the availability of food has increased dramatically. This has been paralleled with the increase in obesity rates. In 2015 almost 2 billion people were estimated to be overweight (Chooi et al., 2019). Obesity predisposes several illnesses such as cancer, type 2 diabetes, heart disease, stroke, and mental illnesses such as depression (Seabrook & Borgland, 2020). Obesity results from positive energy imbalance and recent studies have focused on the role of the central nervous system in metabolic dysregulation. One candidate mechanism behind obesity is the altered function of the brain’s reward circuit and dysfunction in volitional control of appetite (Nummenmaa, Hirvonen, et al., 2012; Tuulari et al., 2015). The imbalance between the prefrontal control mechanism and the striatal reward circuits generating motivational signals upon encountering food may thus lead some individuals to overeat despite their current metabolic status (Drelich-Zbroja et al., 2022). Obese subjects have elevated striatal metabolism which is also linked with amplified reward responses to appetizing foods (Nummenmaa, Hirvonen, et al., 2012). Moreover, body-mass index (BMI) is also positively associated with activation of the taste cortices while tasting sweet solutions, indicating sensory preference for high-calorie foods (Chen & Zeffiro, 2020).

Functional MRI (fMRI) studies have established that premotor areas, superior frontal cortices and the precuneus regulate cognitive control of appetite while viewing food cues (Tuulari et al., 2015). These areas play key roles in the brain’s cognitive inhibition network (Laird et al., 2011; Liddle et al., 2001; Tuulari et al., 2015). In turn, feeling hungry have been associated to increased activation of insula, thalamus and parahippocampal gyrus (Zhao et al., 2017). Compared to normal-weight individuals, obese individuals had lowered responses in dorsal striatum during volitional appetite control, while normal-weight individuals had stronger activations in bilateral dorsal caudate nuclei (Tuulari et al., 2015). In obese subjects, reduced activity has also been found in other components of the inhibitory control system, such as in the supplementary motor area (SMA) (Chen & Zeffiro, 2020). The activity of the dorsolateral prefrontal cortex (DLPFC), a key node in the brain’s inhibitory network governing food intake, is dampened in obese versus normal-weight individuals (Gluck et al., 2017). In addition, increased activity of DLPFC has been observed to predicts healthier food choices and better dietary restraint (Parsons et al., 2021; Zhao et al., 2017). In line with this, dysfunction of DLPFC has been observed in several mental health disorders such as binge eating disorder and substance use disorders (Gluck et al., 2017).

Sociability is often considered as the “default mode” of human brain function, given the centrality of social interaction to our species (Hari et al., 2015). Interestingly, recent neuroimaging work also highlights that subset of the brain regions involved in social perception are also activated when seeing others eating, highlighting the intertwined nature of food and sociability in the brain (Santavirta et al., 2023). We understand others partially by “copying” their behaviours and internal states in our own minds. There is ample evidence of such embodied vicarious representation of others motor, motivational and affective states (Katsyri et al., 2013; Mobbs et al., 2009; Nummenmaa, Glerean, et al., 2012; Rizzolatti & Craighero, 2004; Singer et al., 2004). Together with the data on the tendency to overeat in the presence of others (Higgs & Thomas, 2016; Ruddock et al., 2021) these data suggest that the tendency to automatically remap others’ feeding behaviour in the observers’ brain could be a potent modulator of feeding and food-induced reward. However, this hypothesis currently lacks empirical support.

### The current study

Here we measured haemodynamic brain responses to naturalistic episodes of social eating in short movie scenes and correlated the strength of the responses with subjects’ BMI. We hypothesized that watching social eating would result in a vicarious feeding response, manifested in increased somatomotor and affective engagement in the brain. Because previous studies have also linked dysfunctional inhibitory control systems with obesity, we also predicted that the participants’ BMI would modulate the brain responses for social feeding in brain areas linked with volitional inhibitory control, such as in the prefrontal cortex and striatum.

## Methods

### Subjects

A total of 104 healthy volunteers were studied. In addition of the standard fMRI exclusion criteria, we excluded subjects with earlier psychological or neurological disorder, current substant or alcohol abuse and medications that affected to the central nervous system. Two subjects were excluded from further analyses because unusable MRI data due to gradient coil malfunction and two subjects were excluded because of anatomical abnormalities in structural MRI. Finally, three subjects were excluded due to visible motion artefacts in preprocessed functional neuroimaging data. This yielded a final sample of 97 subjects (50 females, mean age of 31 years, range 20 - 57 years, BMI range 18.2- 30.8, mean 22.5, SD 3.54). All subjects gave an informed, written consent and were compensated for their participation. The study protocol was approved by the ethics board of the Hospital District of Southwest Finland and the study followed the Declaration of Helsinki.

### Stimulus

To map brain regions that are activated while viewing eating, subjects were scanned in fMRI while they were shown short videoclips (median duration 11.2 seconds, range 5.3 – 28.2 seconds, total duration 19 min 44 seconds). Movies were shown consecutively without breaks in fixed order for all participants. The clips were selected from various Hollywood movies, and they showed humans in different everyday situations (e.g eating, talking, sleeping, interacting etc.). Five independent annotators rated the moment-to-moment presence and magnitude of eating from the stimulus film clips and the regressor for eating was calculated as average over the annotators. To extract the eating related heamodynamic responses from other social information processing related to observing films, the brain responses to eating were contrasted with those of seeing people standing (People did not eat while they were standing in the stimulus films). See Figure 1 for the time series of the presence of eating and standing. Visual stimuli were presented with NordicNeuroLab VisualSystem binocular display. Sound was conveyed with Sensimetrics S14 insert earphones. Stimulation was controlled with Presentation software. Before the functional run, sound intensity was adjusted for each subject so that it could be heard over the gradient noise.

**Figure 1.**
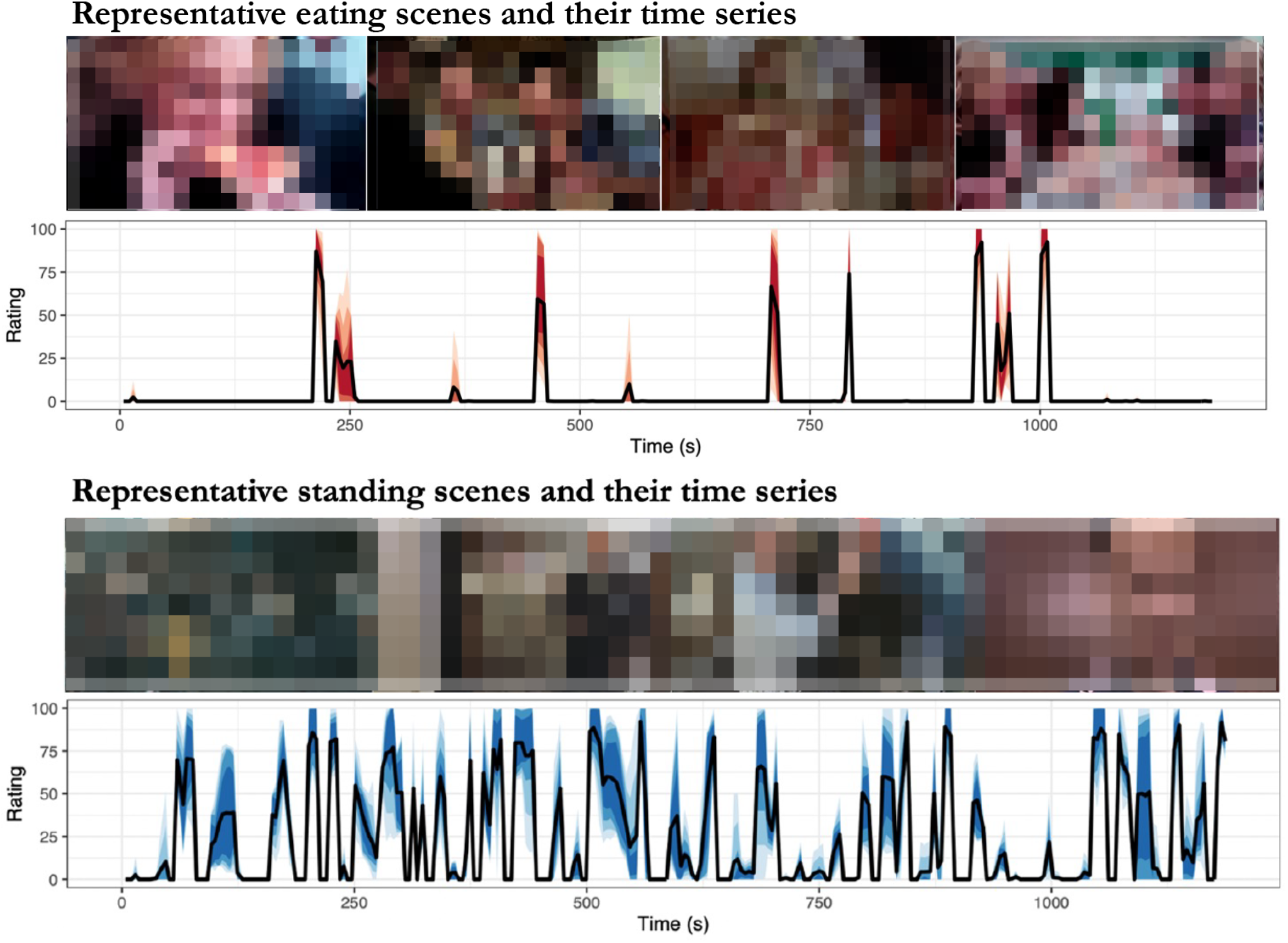
Representative eating (top row) and standing (bottom row) scenes with the corresponding intensity time series of the events.

### MRI data acquisition

The MRI data were acquired using a Phillips Ingenuity TF PET/MR 3-T whole-body scanner. High-resolution (1 mm^3^) structural images were obtained with a T1-weighted sequence (TR 9.8 ms, TE 4.6 ms, flip angle 7°, 250 mm FOV, 256×256 reconstruction matrix). Functional images were obtained for the movie experiments, respectively, with a T2*-weighted blood-oxygenation-level-dependent (BOLD) echo-planar imaging sequence (TR 2600 ms, TE 30 ms, 75◦ flip angle, 240 mm FOV, 80×80 reconstruction matrix, 62.5 kHz bandwidth, 3.0 mm slice thickness, 45 interleaved slices acquired in ascending order without gaps).

### MRI data preprocessing

MRI data were preprocessed using fMRIPprep 1.3.0.2 (Esteban et al., 2019). The following preprocessing was performed on the anatomical T1-weighted (T1w) reference image: correction for intensity non-uniformity, skull-stripping, brain surface reconstruction, and spatial normalization to the ICBM 152 Nonlinear Asymmetrical template version 2009c (Fonov et al., 2009) using nonlinear registration with antsRegistration (ANTs 2.2.0) and brain tissue segmentation. The following preprocessing was performed on the functional data: coregistration to the T1w reference, slice-time correction, spatial smoothing with a 6-mm Gaussian kernel, non-aggressive automatic removal of motion artifacts using ICA-AROMA (Pruim et al., 2015), and resampling of the MNI152NLin2009cAsym standard space. Low-frequency drifts were removed with a 240-s-Savitzky– Golay filter (Çukur et al., 2013).

### Full-volume GLM data analysis

The fMRI data were analyzed in SPM12 (Welcome Trust Center for Imaging, London, UK, http://www.fil.ion.ucl.ac.uk/spm). To reveal regions activated by eating and standing, a general linear model (GLM) was fitted to each subject’s voxelwise BOLD-signals separately. The first-level fixed effects model included regressors for eating and standing and eight low-level audiovisual features and signals from cerebrospinal fluid and white matter as confounds. We used our previously validated low-level model for controlling the potential low-level audiovisual confounds in the movie clips (Santavirta et al., 2023). Briefly, 14 audiovisual features were extracted from the movie clips and principal component analysis (PCA) revealed that eight principal components explained over 90% of the total variance of the audiovisual features. These eight principal components were included in the first-level model. All regressors were convolved with canonical double-gamma HRF before analyses. First-level contrast images were then defined for the main effects of eating and standing as well as for the subtraction between eating and standing (eating – standing). Finally, each participant’s contrast images were subjected to a second level analysis. In the second level, we modelled the association between the participants’ BMI to the BOLD responses for eating and standing. The second level model included participants’ BMI, age, and sex. The second level results were statistically tested using parametric one-sample t-tests.

### Region-of-interest analyses

To visualize the results, weights for viewing eating and standing were extracted in bilateral masks defined by ROIs extracted from AAL2 atlas (Rolls et al., 2015) added with more fine-grained parcellations for precentral gyrus, postcentral gyrus and nucleus accumbens from Brainnetome atlas (Fan et al., 2016). The mean beta weights for each ROI were calculated from each subject’s contrast images and the second level modeling was conducted similarly with the full volume analysis.

## Results

### Regional responses to vicarious feeding

Across all subjects viewing eating increased BOLD activity in primary motor and premotor cortex, temporal cortex, somatosensory cortex, thalamus, and parahippocampal gyrus (Figure 2). Eating related brain responses were significantly higher compared to the responses for standing in primary motor and premotor cortex, somatosensory cortex, SMA, posterior parietal cortex, visual cortex, DLPFC, insula, thalamus, para hippocampal, middle temporal and superior occipital gyrus and precuneus (Figure 3).

**Figure 2.**
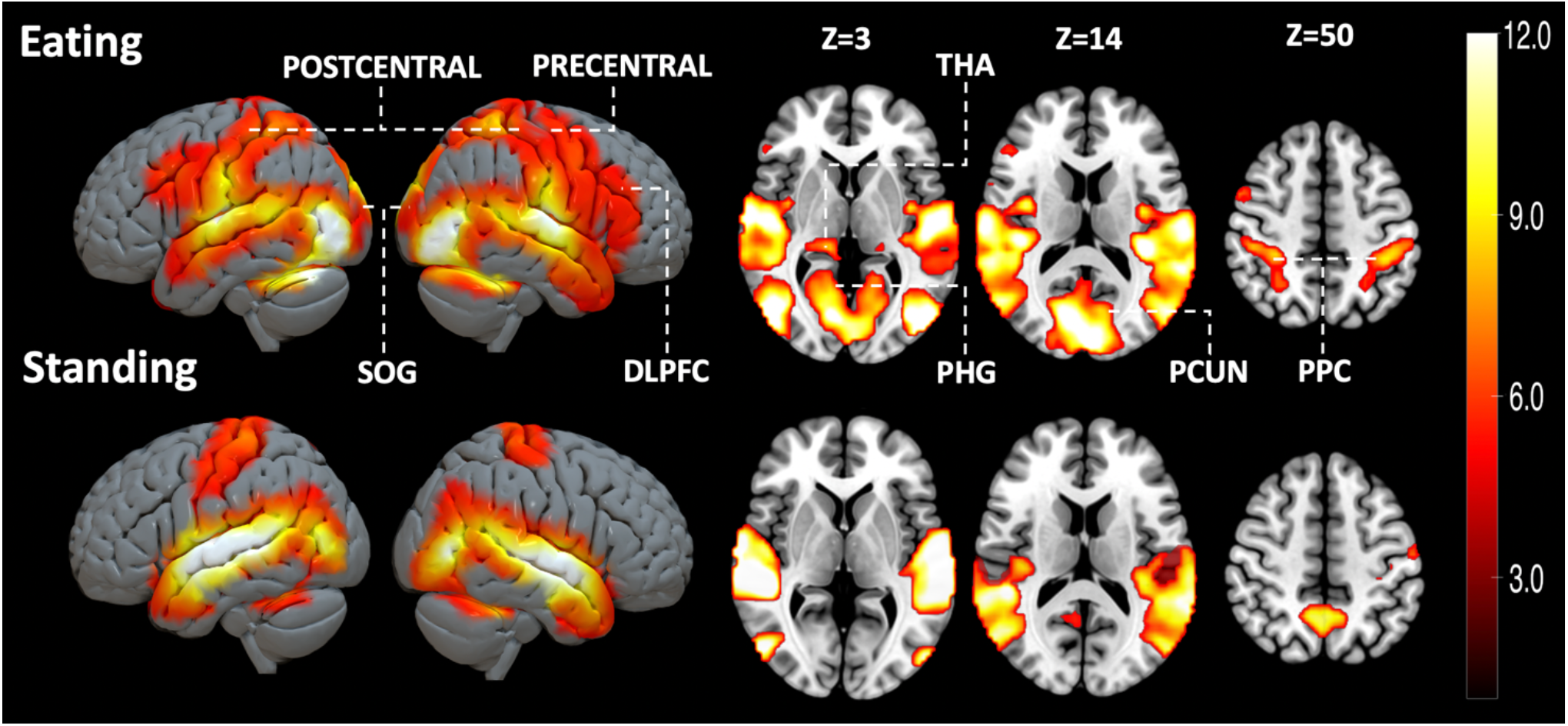
Brain responses for viewing feeding and standing in social scenes (FWE-corrected on the voxel level, alpha=0.05). DLPFC=Dorsolateral prefrontal cortex, PCUN=Precuneus, PHG=Parahippocampal gyrus, PCUN=Precuneus, PRECENTRAL=Precentral gyrus, POSTCENTRAL=Postcentral gyrus, PPC=Posterior parietal cortex, SOG=Superior Occipital Gyrus, THA=Thalamus.

**Figure 3.**
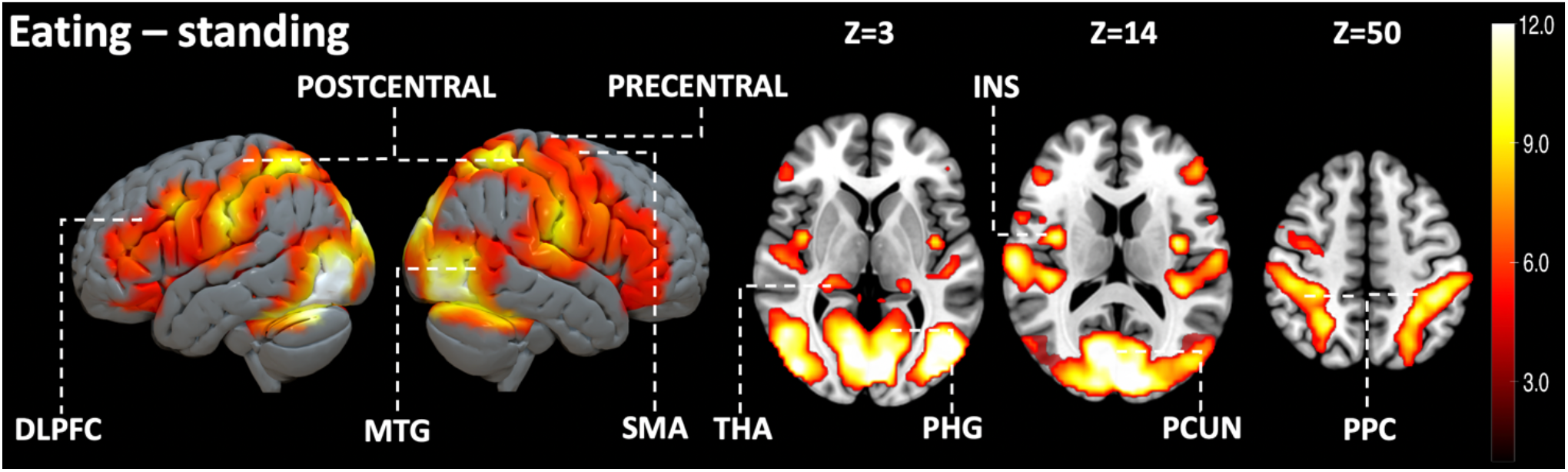
Brain regions responding more strongly to perceived eating than standing (FWE-corrected on the voxel level, alpha=0.05). DLPFC=Dorsolateral prefrontal cortex, INS=Insula, MTG=Middle temporal gyrus, PCUN=Precuneus, PHG=Parahippocampal gyrus, PRECENTRAL=Precentral gyrus, POSTCENTRAL=Postcentral gyrus, PPC=Posterior parietal cortex, SMA=Supplementary motor area, THA=Thalamus.

### BMI-dependent responses to viewing feeding

We next tested whether the responses to vicarious feeding would be associated with subjects’ BMI. This analysis revealed that BMI was negatively associated with eating-evoked BOLD-signals in DLPFC, primary motor cortex, precuneus, parahippocampal gyrus, thalamus, putamen and caudate nuclei (Figure 4). While eating was associated with stronger BOLD responses than standing in various brain regions this difference in BOLD response became weaker with increasing BMI (negative association between BMI and eating – standing contrast) in DLPFC, primary motor cortex, precuneus, parahippocampal gyrus, putamen, and caudate nuclei (Figure 5). Scatterplots in figure 6 show the regional negative associations between brain responses to perceived eating and BMI. Finally, scatterplots in figure 7 show the regions where the difference in brain responses for eating and standing diminished with increasing BMI.

**Figure 4:**
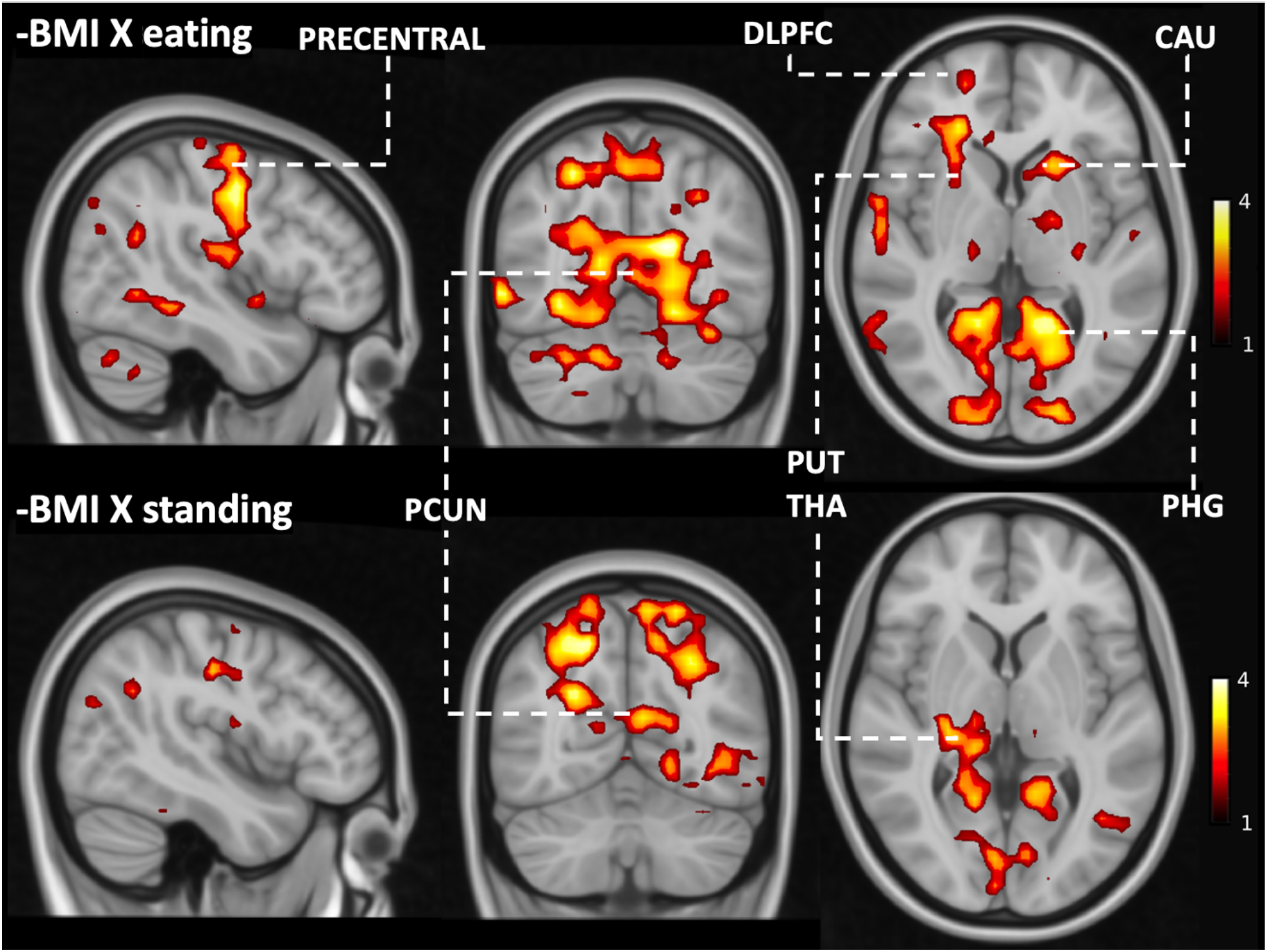
Brain regions where BMI was negatively associated with viewing eating and standing (FWE-corrected on the cluster level, cluster forming threshold: p < 0.05). DLPFC=Dorsolateral prefrontal cortex, CAU=Caudate Nuclei, PHG=Parahippocampal gyrus, PRECENTRAL=Precentral gyrus, PUT=Putamen, THA=Thalamus.

**Figure 5.**
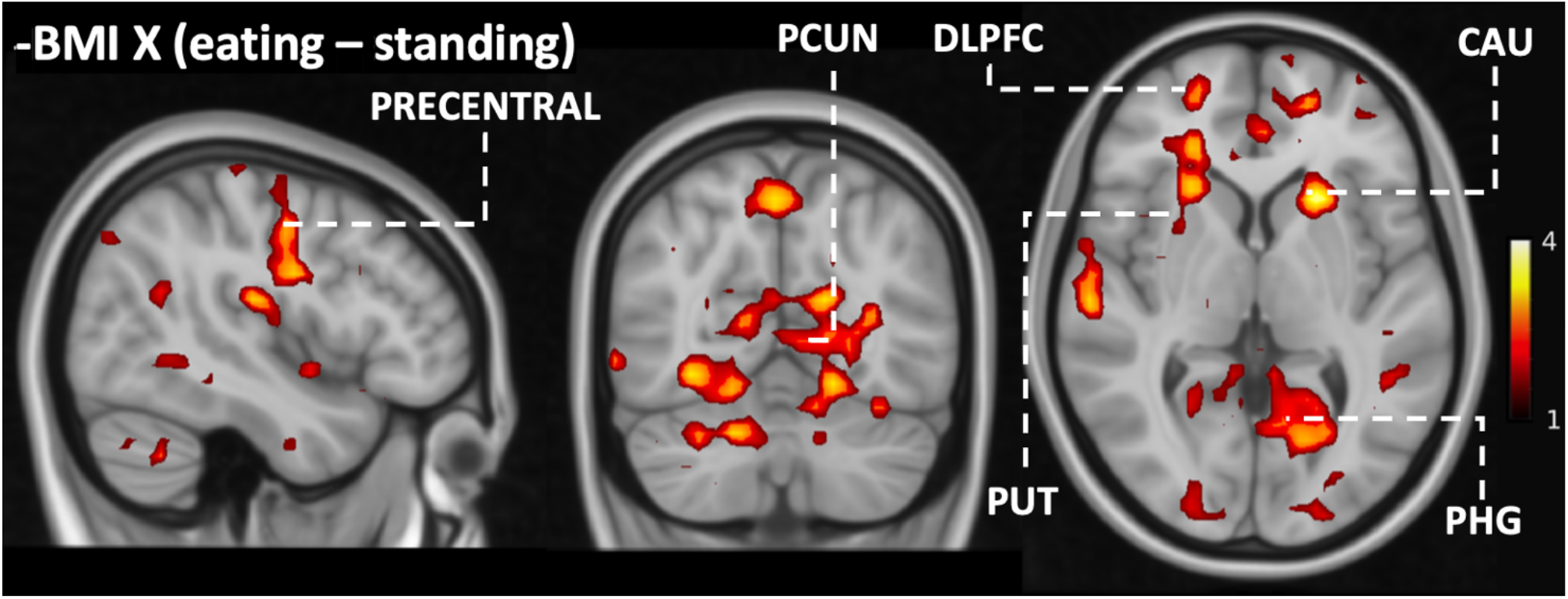
Brain regions where BMI was negatively associated with viewing eating versus standing (FWE-corrected on the cluster level, cluster forming threshold: p < 0.05). DLPFC=Dorsolateral prefrontal cortex, CAU=Caudate nuclei, PCUN=Precuneus, PHG=Parahippocampal gyrus, Precentral=Precentral gyrus, Put=Putamen.

**Figure 6:**
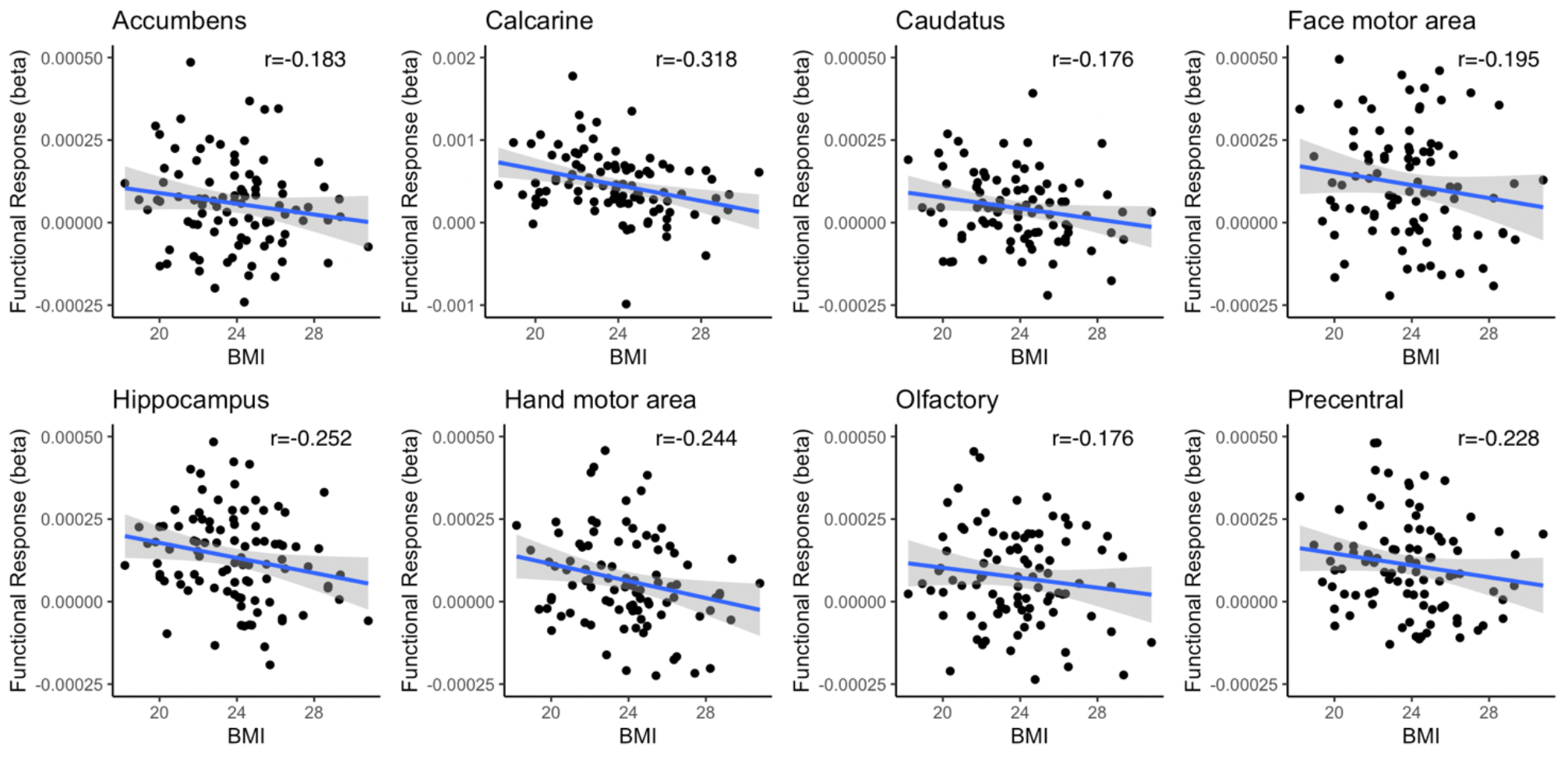
Regional associations between BMI and haemodynamic responses to vicarious eating in representative ROIs. Note that the scatterplots are used for visualization and the statistical inference is based on the full-volume analysis.

**Figure 7.**
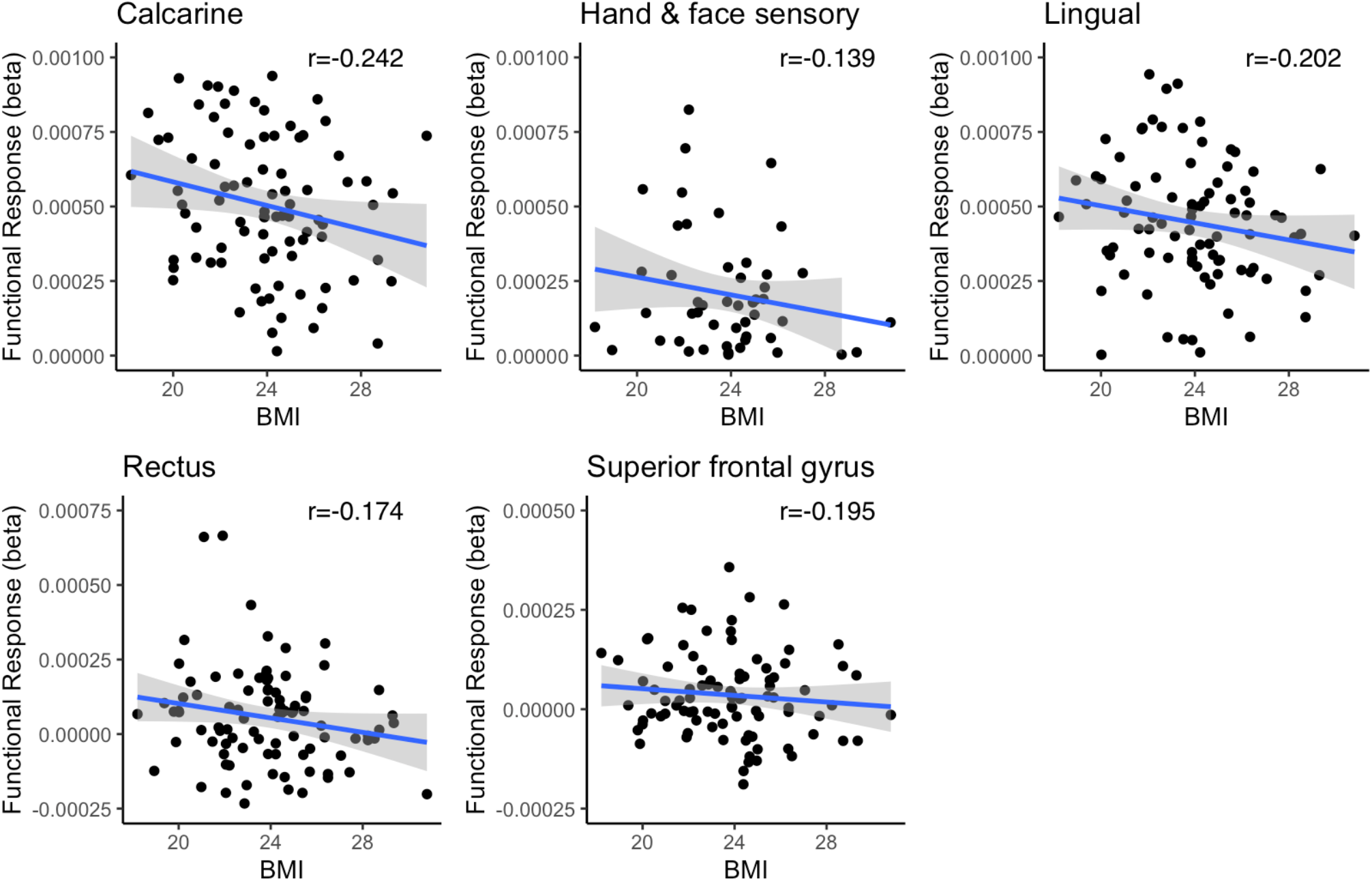
Regional associations between BMI and haemodynamic responses to vicarious eating versus standing in representative ROIs (p<0.05). Note that the scatterplots are used for visualization and the statistical inference is based on the full-volume analysis.

## Discussion

Our main finding was that watching eating activates brain areas subserving voluntary movements such as premotor cortex, primary somatosensory cortex, somatosensory association area, SMA and DLPFC but also the areas that are linked with sensation of hunger such as thalamus and insula (Bhattacharjee et al., 2021; Gluck et al., 2017; Ryun et al., 2023; Zhao et al., 2017). Additionally, we found that the vicarious feeding responses in the brain were negatively associated with subjects’ BMI, such that higher BMIs were linked with weaker responses. All in all, our results show that the human brain continuously “mirrors” others’ feeding behaviours potentially to promote social feeding, and that this process is downregulated in individuals with high BMI.

### Brain responses for vicarious eating

Across all subjects, vicarious eating activated precentral and postcentral gyrus, premotor cortex, DLPFC, somatosensory association area, thalamus, and insula. Primary motor cortex in precentral gyrus controls volitional muscle motions whereas premotor cortex organizes complex movements with cognitive functions (Bhattacharjee et al., 2021). DLPFC and insula are both associated with cognitive control of eating and appetite control (Gluck et al., 2017; Tuulari et al., 2015). In turn, somatosensory cortices are centrally involved in tactile perception but also in emotional perception and simulating others’ mental states (Nummenmaa et al., 2014). Thalamus in turn contributes to a multitude of affective processes including arousal modulation (Laird et al., 2011). Direct comparison between viewing eating versus standing revealed increased activation in precentral and postcentral gyrus, DLPFC, posterior parietal cortex, MTG, PHG, thalamus and insula stronger than perceiving people standing. In previous studies precentral and postcentral gyri have been linked with disinhibition to eat (Zhao et al., 2017). Hunger has been associated to increased activity of insula, right thalamus and PHG (Zhao et al., 2017). DLPFC participates in volitional appetite control (Tuulari et al., 2015). In turn posterior parietal cortex has been associated to participate in decision making and motor function (Leoné et al., 2014; Lindner et al., 2010).

Overall, our results suggest that the brain regions participating in voluntary movements, somatosensation and reward processing activate during vicarious eating, likely reflecting mental simulation of the actions and emotions associated with first-hand feeding similarly as has previously been established for emotions and various motor actions states (Katsyri et al., 2013; Nummenmaa, Glerean, et al., 2012; Rizzolatti & Craighero, 2004; Singer et al., 2004). We propose that this tendency to internally mimic others feeding in social contexts might be a powerful cue for increasing appetite and initiating feeding. Watching eating results in somatomotor and affective “mirroring” response of actual feeding in the brain, which may at least partly explain why people tend to eat more together than alone (Higgs & Thomas, 2016; Ruddock et al., 2021). The visceral and affective engagement could trigger an anticipatory reward responses engaging an urge to eat independently of the current metabolic state, potentially increasing the rewarding value of foods when eating in the company of others. (Huang et al., 2022). Finally, the automatic motor preparation of feeding-related actions seen in others could lower the threshold for engaging in feeding.

### BMI-dependent responses for vicarious eating

Our second main finding was that the neural responses to vicarious feeding were modulated by BMI. Specifically, responses to viewing feeding versus standing were negatively correlated with BMI in caudate nuclei, putamen, primary motor cortex and parahippocampal gyrus (PHG). Of these regions, the primary motor cortex enables voluntary movement. (Bhattacharjee et al., 2021). PHG participates in satiety control (Brooks et al., n.d.). Putamen and caudate nuclei in turn participate in motor inhibition and processing (Chen & Zeffiro, 2020; Tuulari et al., 2015). Importantly, the BMI-dependent variation in the motor strip was observed specifically in the face and hand areas (Fig 6), suggesting that the effect directly pertains with feeding-related actions.

The striatum and particularly the caudate nuclei are important components of the human reward circuit, and unexpectedly they were not significantly activated in the primary analysis contrasting viewing eating versus standing. However, we found that the striatal activations were dependent on the subjects’ BMI. The larger the BMI, the weaker the striatal responses were. This indicates that the striatal reward encoding of vicarious eating is downregulated in obesity. In line with this, experimental studies have indeed found that when eating alone, overweight children eat more than normal-weight children, but this difference is abolished when eating in a group (Salvy et al., 2007). Similarly, obese adults eat very little when in the company of lean individuals (such as those in our stimuli) whereas their food consumption is significantly amplified when feeding with an obese individual (de Luca & Spigelman, 1979). Taken together these results suggests that obesity and overweight might be associated with different social norms regarding feeding that may make joint meals less appealing, which would then lead to lowered vicarious feeding responses in the reward circuit. Accordingly, eating together might initially promote obesity, but it is possible that this trend is subsequently curbed following weight gain. However, our cross-sectional study cannot directly address this issue.

Finally, BMI-dependent variation in the vicarious feeding responses were also observed in PHG and DLPFC. PHG has been discovered to participate in satiety control (Brooks et al., n.d.) while DLPFC participates in cognitive control, regulates food intake via cognitive appetite regulation (Gluck et al., 2017). Accordingly, modulation of the DLPFC and PHG activity by BMI might reflect aberrant inhibitory control over visually induced appetite. In sum, the BMI-dependent alterations in the vicarious feeding responses likely highlight three distinct processes: lowered tendency for motor simulation, lesser affective engagement, and lower engagement of fronto-cortical control circuits. Whereas the two first processes might make high-BMI individuals less likely to eat when with others due to lowered affective and motor impulses, the dampened DLPFC activation might partially counteract the lowered affective and motor impulses. This hypothesis however needs to be validated in future studies.

### Limitations

The BMI range of our subjects was relatively narrow and there was only one obese subject in our study. Most of our subjects were either normal weight or overweight individuals. Hence, our results mainly pertain with BMI-dependent modulation of vicarious feeding responses in predominantly subjects. The foods shown in the stimulus scenes were both palatable and nonpalatable. Therefore, unlike most fMRI studies with pictorial food stimuli, our results do not distinguish the brain activation patterns for reward-dependent encoding of foods. Finally, the video viewing protocol did not allow strict control over the audiovisual features of the different stimulus categories because we wanted to focus on natural dynamic episodes representative of real-life social eating. We however performed extensive statistical control for the sensory features and the effects remained significant even after such controls.

## Conclusions

We conclude that vicarious eating activates brain regions that participate in voluntary movements and process sensory information. This affective and somatomotor “mirroring” of the emotional and motor components of food intake might prepare the observer for joining the meal thus promoting food intake. These responses were however dampened as a function of the BMI of the subjects. Our results thus demonstrate the importance of the social context of eating and show how visual representations of others’ feeding are transformed into somatomotor and affective representations possibly promoting appetite and feeding. Future studies need to elucidate how these vicarious feeding responses contribute to actual food intake and development of obesity.

## Acknowledgement

The study was supported by the Sigrid Juselius Foundation and Academy of Finland (grants numbers 294897 and 332225) to LN, Turku University Foundation and Alfred Kordelin Foundation grants to SS and Finnish Governmental Research Funding for Turku University Hospital and for the Western Finland collaborative area to SS.

